# Working memory-related activity in catecholaminergic nuclei in schizophrenia

**DOI:** 10.1101/2023.08.28.555235

**Authors:** Nada Amekran, Verónica Mäki-Marttunen

## Abstract

Schizophrenia is a complex psychiatric condition in which cortical, subcortical and neuromodulatory alterations have been implicated in its symptom expression. Long standing views of schizophrenia symptoms have posed that alterations in catecholaminergic systems, which explain psychotic symptoms, may be also associated with the cognitive impairments commonly observed in this condition. However, evidence on the involvement of catecholaminergic regions on cognitive functions in schizophrenia remains scarce. Working memory is one cognitive domain where schizophrenia patients present more impairments at higher levels of cognitive load. Here we explored the activation of catecholaminergic regions during a working memory task in schizophrenia. We reanalyzed an openly available functional magnetic resonance imaging dataset where schizophrenia patients and healthy controls were scanned while performing the N-back task. We compared activation of two dopaminergic areas, ventral tegmental area and substantia nigra, and of a noradrenergic nucleus, locus coeruleus, to the presentation of targets, and compared three different levels of cognitive load (0-, 1– and 2-back). We found that across nuclei, higher load was related to lower activation. Furthermore, schizophrenia patients showed reduced activation at the highest load level when compared to healthy controls. These findings point to catecholaminergic systems as mediators of the deficits in effort processing in schizophrenia. Our study lends further support for the importance of including catecholaminergic systems in the mechanisms of cognitive deficits in schizophrenia.

## Introduction

Schizophrenia is a psychiatric disorder mainly characterized by positive (e.g. hallucinations or delusions), negative (e.g. reduced emotion expression or decreased motivation), and cognitive (e.g. decreased executive functioning) symptoms (Cardno & Gottesman, 2000; McGrath et al., 2008; Owen et al., 2016; Sullivan et al., 2003). As part of the cognitive symptoms, deficits in working memory (WM) are commonly observed (Barch & Smith, 2008; Kalkstein et al., 2010; Lee & Park, 2005). Several attempts to explain the mechanisms of the complex symptomatology of schizophrenia led scientists to investigate the function of the different neurotransmitter and neuromodulatory systems in schizophrenia (Stahl, 2018). Despite much advancement in this direction, cognitive deficits remain elusive to treat pharmacologically, and further investigation on the mechanisms behind the expression of cognitive deficits, and the involvement of neuromodulatory systems, is thus warranted.

Currently, there is a general recognition that catecholaminergic systems play a role in schizophrenia symptomatology (Howes et al., 2023; Mäki-Marttunen et al., 2020; Os & Kapur, 2009; Abi-Dargham et al., 2000; Perez-Costas et al., 2010; Tost et al., 2010; Chen et al., 2021). This view is supported by post-mortem studies on brainstem nuclei, structural studies using magnetic resonance imaging (MRI), and pharmacological studies (Rice et al., 2016; Howes et al., 2023; Schulz et al., 2022; Weinstein et al., 2017; van Kammen & Kelley, 1991). However, few studies aimed to directly measure activity in catecholaminergic structures in patients in vivo. One reason is the difficulty in assessing activation in small, deep structures of the brain (Brooks et al., 2013; Sclocco et al., 2018). Recent advances in the methodology to assess activity in brainstem regions using functional MRI (fMRI) and the availability of atlases to assist localization are promising to assess questions regarding the role of catecholaminergic systems in psychiatric disease (Beissner et al., 2014; Matt et al., 2019; Mäki-Marttunen & Espeseth, 2021; Turker et al., 2021). Only a few studies have assessed functional measures of activity in catecholaminergic areas, e.g. ventral tegmental area (VTA), substantia nigra (SN) and locus coeruleus (LC), during different cognitive tasks in schizophrenia (Rausch et al., 2014; Köhler et al., 2019; Cuervo-Lombard et al., 2012). These studies reported a diminished activation of dopaminergic areas VTA and SN, and a possible alteration of locus coeruleus activity. Importantly, how different levels of cognitive load affect the recruitment of those systems has never been assessed.

A common finding in behavioral studies using executive tasks is that schizophrenia patients show relatively unaffected performance at low cognitive load levels, while they perform worse as load increases, even at levels that do not compromise performance in healthy controls (Gjerde, 1983; Granholm et al. 1997). This suggests that basic sensory processing is spared in schizophrenia but patients experience an overload at lower levels of cognitive demand that healthy individuals do not experience. For instance, the N-back task is a working memory task that shows robust differences in performance between healthy participants and schizophrenia patients at higher load levels but not at low load (Carter et al., 1998; Deserno et al., 2012; Starc et al., 2017). This task has also been associated with different levels of brain activation between schizophrenia patients and healthy controls in areas related to working memory processing (Carter et al., 1998; Brahmbhatt et al., 2006; Jiang et al., 2015; Gallucci et al., 2022). Given that the catecholaminergic neuromodulatory systems facilitate effortful behavior (Aston-Jones and Cohen, 2005) and working memory functions (Motley, 2018), a pending question is whether their recruitment is also impaired in schizophrenia patients at different levels of WM load. Answering this question would provide a better overview of how neuromodulatory systems participate in dealing with varying cognitive demands to working memory in schizophrenia.

The goal of the current study is to assess WM-related activity in catecholaminergic structures in healthy individuals as compared to individuals with schizophrenia at different WM load levels. We hypothesized that the activity in these structures would reflect an impaired ability to recruit effort-related modulators in schizophrenia. Alternatively, an absence of difference in activity in these structures would indicate that modulatory activity due to load is intact, and downstream processes such as prefrontal retrieval of targets, would be impaired.

## Materials and Methods

### Participants

Participants were recruited and tested by Repovš and Barch (2012) and data were made available on OpenNeuro. The complete dataset consisted of data from patients with schizophrenia, healthy controls, and siblings of both groups. For this study we did not include the data from siblings of patients with schizophrenia. We added an age requirement of minimum 18, and did not include data of participants younger than 18. We also excluded participants without task design information (sub-72) or with reduced number of volumes (sub-81, 1-back). As we did not look at effects on siblings, we pooled the data of healthy controls and their siblings together into one control group to still have a balanced design after exclusion based on age.

### N-back task

The n-back task is a working memory task where participants are presented with a series of letters in each block. The task is to assess whether the shown letter is the same as a pre-specified target letter (0-back) or the same as a previously shown (1-back/2-back) letter in the sequence. In the 1-back condition, the target letter is the same as the one shown in the trial before, and in the 2-back condition the target letter is the same as the one shown 2 trials before. MRI scans were obtained during the n-back task, with separate runs for the 0-back, 1-back, and 2-back condition. All participants did an n-back task in all conditions. A full description of the task and procedure can be found in Repovš and Barch (2012).

### MRI dataset

Data were downloaded from OpenNeuro (https://openneuro.org/datasets/ ds000115/versions/00001). The original paper describing the full dataset can be found on https://www.ncbi.nlm.nih.gov/pmc/articles/PMC3358772/ (Repovš and Barch, 2012). All scans were acquired on a 3T Tim TRIO scanner. For each participant, data consisted of a T1-weighted structural image (TR = 2400 ms, TE= 3.16 ms, FOV = 256 mm, flip angle= 8°, voxel size = 1 mm x 1 mm x 1 mm) and three task-related functional T2* images (TR = 2500 ms, TE= 27 ms, FOV = 256 mm, flip angle = 90°, voxel size = 4 mm x 4 mm x 4 mm). Task-related scans were obtained during the n-back task. Each run consisted of two blocks of either the 0-back, 1-back, or 2-back task, resulting in three 4D volumes for each participant. Each functional scan consisted of 137 volumes. Full details of the data acquisition can be found in the previously mentioned original paper by Repovš and Barch (2012). Data also included event files for each n-back, specifying onset times for each stimulus throughout each functional scan.

### Preprocessing and analysis

Preprocessing was done using the standard pipeline of fmriprep (Esteban et al. 2019). The pipeline included removal of the first 5 volumes to allow for stabilization of magnetic field, slice timing, realignment, artifact correction using AROMA tool, and normalization to MNI152NLin64sym space. This resulted in a preprocessed 4D fMRI image per n-back condition for each participant. The voxel dimensions of the images were 2×2×2 mm^3^. The functional images were submitted to first level analysis using SPM toolbox (https://www.fil.ion.ucl.ac.uk/spm/). For this model, onset times (specified in seconds) and duration of target and non-target stimuli were imported from the original dataset. The canonical hemodynamic response function with its temporal derivatives were used to model each event. We then calculated the contrasts to obtain target-related activation during the 0-back, 1-back and 2-back conditions.

### Brainstem ROI analysis

Regions of interest (ROI) for the catecholaminergic nuclei were obtained from the Brainstem Navigator (Singh et al., 2021; Bianciardi et al., 2015, Figure 1). These nuclei were left and right locus coeruleus (LC), left and right substantia nigra (SN), and left and right ventral tegmental area nucleus complex (VTA). The left and right masks were combined to form one mask per region of interest. Figure 1 shows the masks overlapped on the structural scan of a sample subject. Beta-coefficients within each mask and for each subject and load condition were extracted using the marsbar toolbox (Brett et al., 2002; retrieved from http://marsbar.sourceforge.net).

**Figure 1.**
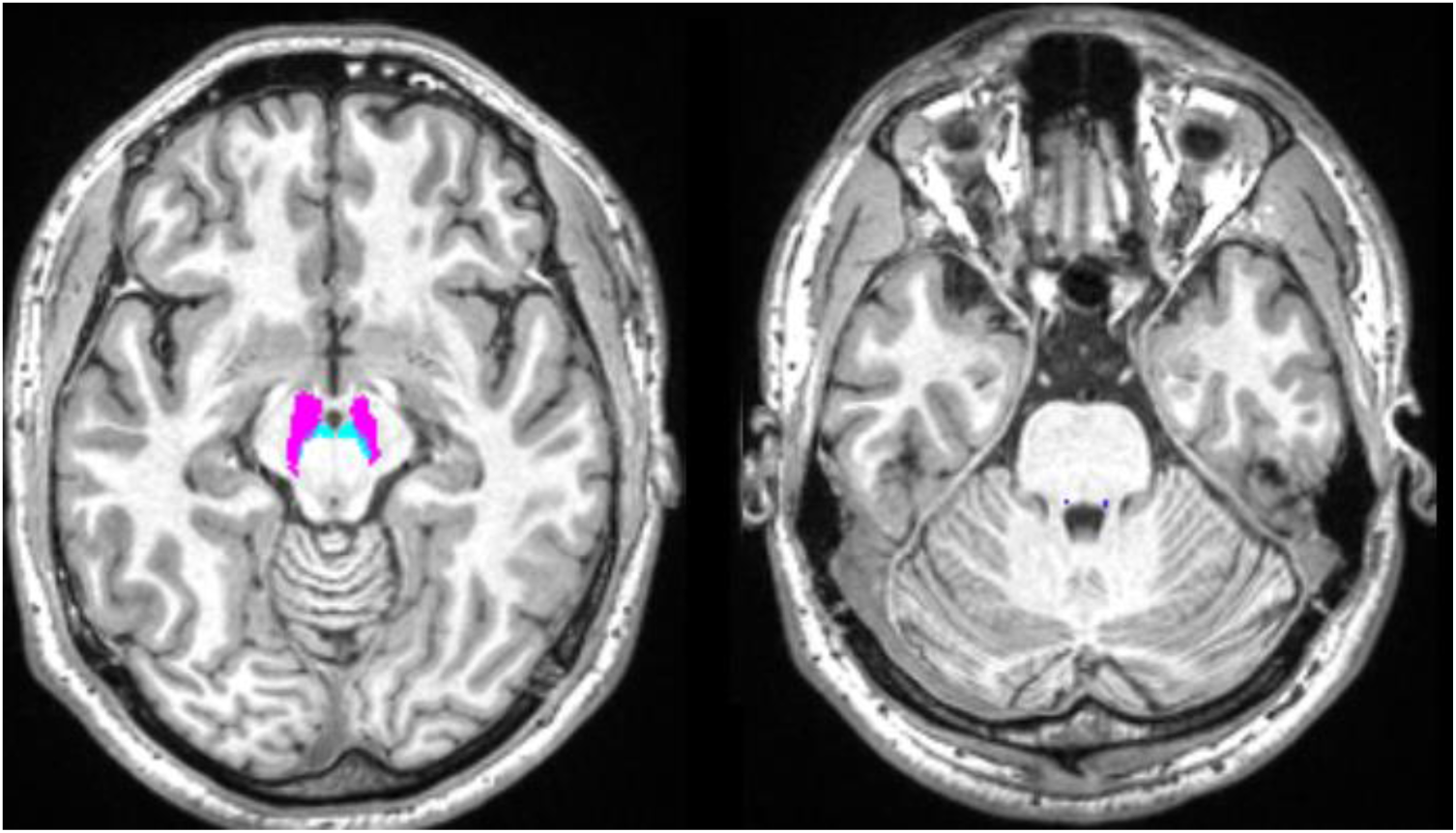
Masks used for ROI analysis overlapped on a sample subject’s T1w volume from fmriprep (sub-01_space-MNI152NLin6Asym_desc-preproc_T1w.nii). VTA (cyan), SN (violet) and LC (blue).

### Statistical analysis

Repeated measures analyses of variance (rmANOVA) were conducted separately for accuracy and reaction time on the n-back task. Factors of interest were group (control or patient group) as between-subjects factor and cognitive load (0-back, 1-back, 2-back) as within-subjects factor. An rmANOVA was conducted on the beta coefficients with group (control or patient group) as between-subjects factor and ROI (LC, SN, VTA) and load (0-back, 1-back, 2-back) as within-subjects factor. fitrm and anova functions as implemented in MATLAB were used to define the linear models and obtain the statistical values. All the codes used for this study are available on github (https://github.com/VeronicaMaki-Marttunen).

### Data Availability

All data has been made available in OpenNeuro (https://openneuro.org/datasets/ ds000115/versions/00001) by the authors of the original work (Repovš and Barch, 2012).

## Results

### Demographics

A total of 28 healthy controls and 22 patients with schizophrenia were included in the final sample. The subject codes are reported in Supplementary table 1. The two groups did not differ significantly in age, while IQ score and years of school were significantly lower in schizophrenia (Table 1).

**Table 1.**
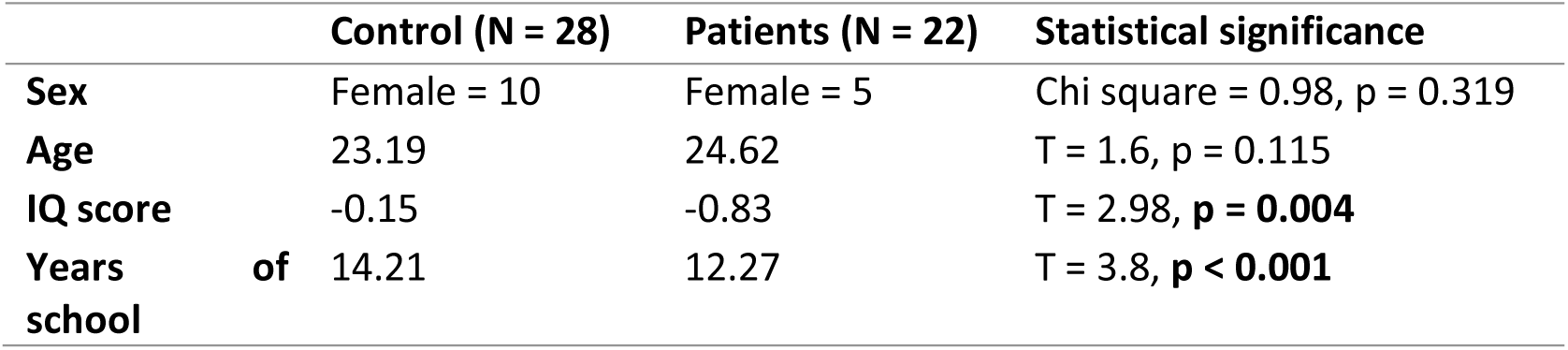
Means (standard deviations) of demographic differences on gender, age, standardized IQ score, and years of school attended. Significant differences were tested at p < 0.05.

### Behavior

We first compared the accuracy and reaction times (RT) between healthy controls and schizophrenia patients. As expected, we found a significant effect of load on accuracy and RT (Figure 2, Table 2), with lower accuracy and higher RTs with higher load. We found a significant effect of group only on accuracy, with lower overall accuracy in schizophrenia patients. We also found a significant group by load interaction, where accuracy was significantly lower in schizophrenia patients but only for higher load levels (two-samples T tes: 0-back: p = 0.267; 1-back: p = 0.003, 2-back: p < 0.001).

**Figure 2.**
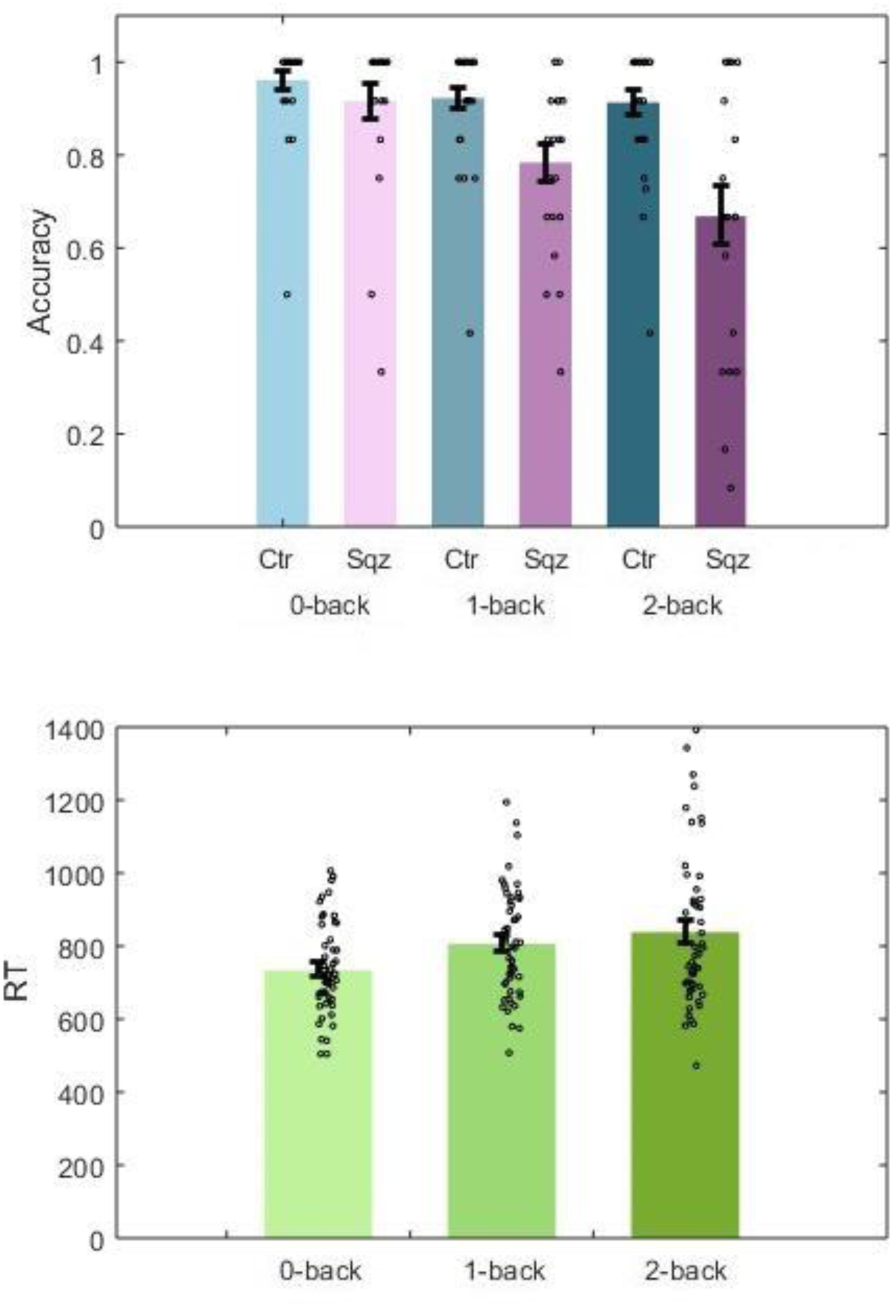
Top: Estimated marginal means of accuracy on each n-back level and group. Bottom: Estimated marginal means of RT on each n-back level across groups.

**Table 2.**
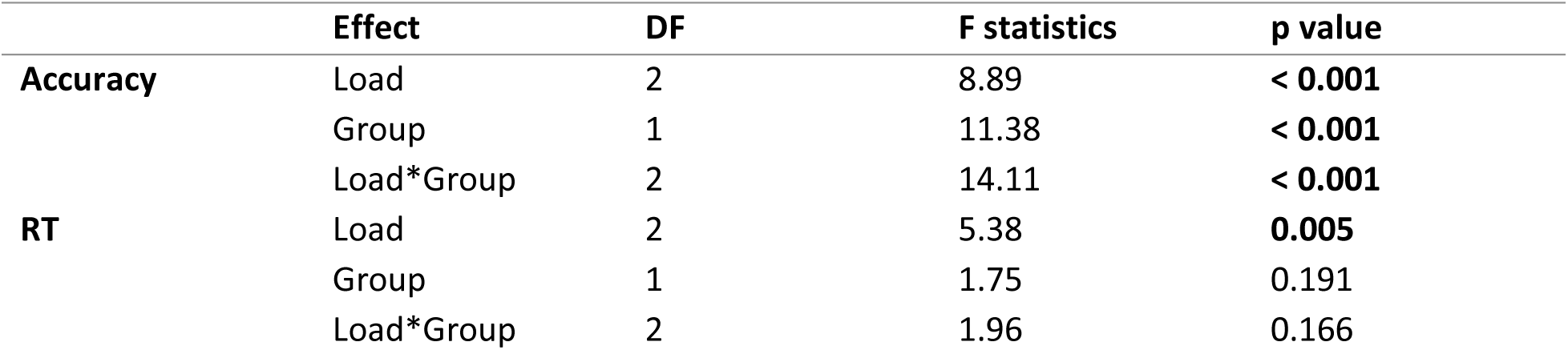
Results of the general linear models applied to Accuracy and RT (see Methods). DF: degrees of freedom.

### Regions of interest analysis

We then analyzed the WM-related activity in three catecholaminergic nuclei: locus coeruleus, substantia nigra, and ventral tegmental area. We found a significant effect of load (Figure 3, Table 3), where the lowest load level evoked significantly larger activity than the higher load levels (paired T test: 0 vs 1 back: p < 0.001; 0 vs 2 back: p = 0.025; 1 vs 2 back: p = 0.066). Overall activation was lower in schizophrenia patients, but this difference was not significant. We also found a significant group by load interaction, where activation was lower in schizophrenia patients compared to healthy controls but only in the highest load level (p = 0.004).

**Figure 3.**
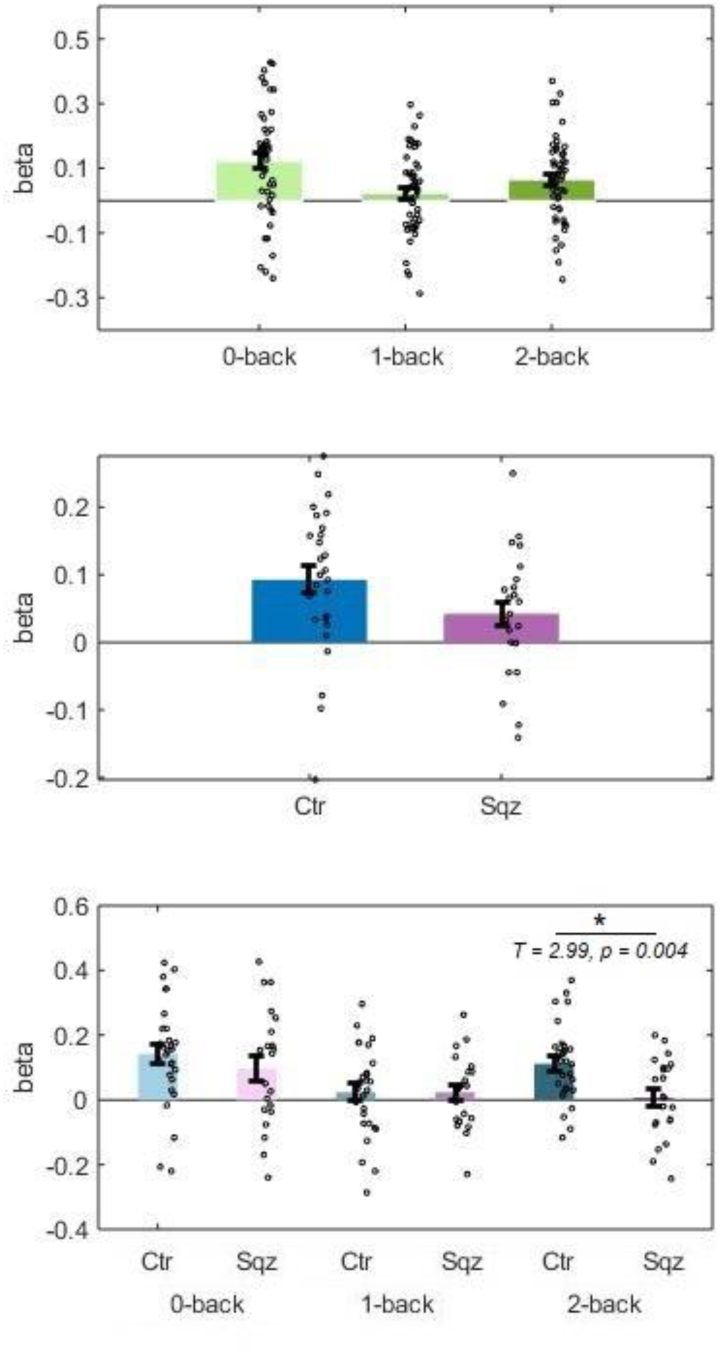
Top: Estimated marginal means of beta coefficients on each n-back level across ROIs. Middle: Estimated marginal means of beta coefficients on each group. Bottom: Estimated marginal means of beta coefficients for each n-back level across group. Asterisk represents significant differences between patients and controls.

**Table 3.**
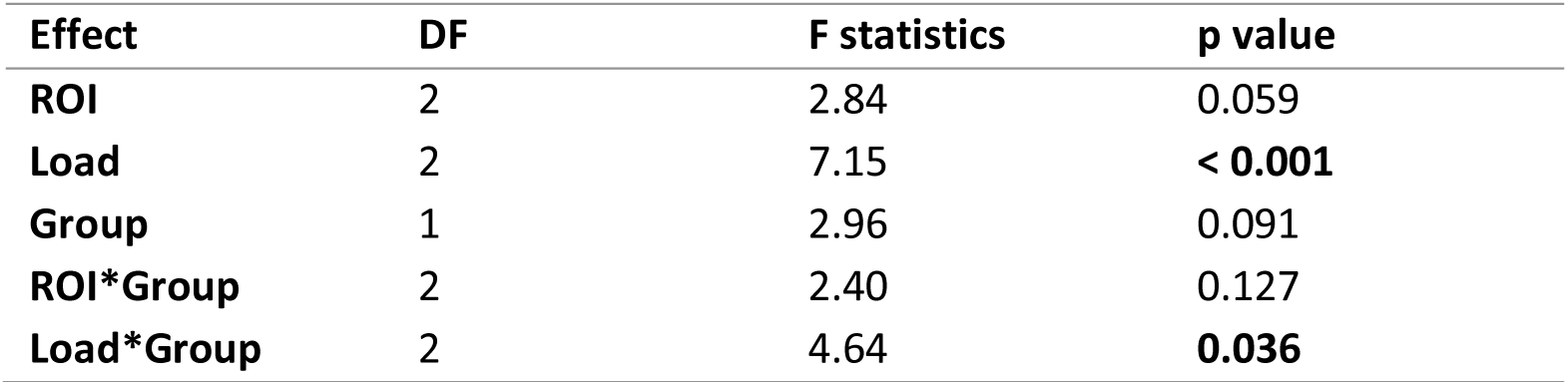
Results of the general linear models applied to the beta coefficients extracted from the regions of interest. DF: degrees of freedom.

## Discussion

In this study, we aimed to assess WM-related activity in catecholaminergic brainstem nuclei in healthy subjects and schizophrenia patients. We compared the activation to a target letter that was presented individually in a stream of letters (0-back, no load), or to targets that were any letter that appeared repeated one place (1-back) or two places (2-back, high load) apart. We found that at the highest load, activity in catecholaminergic nuclei was lower in patients, while there was no difference at lower load levels. This result paralleled the largest reduction in performance at highest load. Our results suggest that a deficient recruitment of catecholaminergic structures may play an important role in the cognitive deficits that characterize the disease.

We studied the activation of the subcortical nuclei VTA and SN (the two main dopaminergic areas of the brain), and of the LC (the main noradrenergic nuclei), during a working memory task. We found that the activity in these nuclei was higher in the control group than the schizophrenia group but only at the highest load level. We also found that the pattern was similar across all regions. The results suggest that catecholaminergic nuclei were successfully recruited in healthy individuals to deal with the higher working memory demands imposed by the higher load level. This is consistent with the role that catecholamines play in facilitating neural activation in demanding cognitive tasks (Aston-Jones & Cohen, 2005; Varazzani et al., 2015), in particular when information must be kept in working memory (Wang et al., 2007; Robbins & Arnsten, 2009). A notable finding was a reduced activity across nuclei in intermediate load (1-back), both in healthy controls and schizophrenia. A possible explanation is that the task of comparing two letters presented immediately after each other (1-back) may have involved a different strategy than the task of comparing every letter with a target letter (0-back), which may still have involved the retrieval of a stimulus held in working memory but that needed not be updated (as in the 2-back condition). For instance, a 1-back task could be solved by a strategy employing reactive rather than proactive control (Maki-Marttunen et al., 2019), and a reactive form of control has been found to be preferred in schizophrenia (Lesh et al., 2013). In sum, individuals with schizophrenia presented a lower activation of catecholaminergic nuclei at the highest load level. This suggests the possibility that the catecholamine system fails to respond in patients with schizophrenia at high cognitive load levels.

A lower recruitment of catecholaminergic areas at the highest load level parallels the linear reduction in accuracy with load level in schizophrenia patients. Previous studies also found reduced accuracy in schizophrenia patients at high cognitive load in the N-back task. We found slower reaction times with increasing load, and this effect was similar in both groups. An impairment in working memory is one aspect of the cognitive symptoms that characterize schizophrenia. It has been hypothesized that low performance at high cognitive load may emerge from patients feeling overloaded at lower cognitive load levels than do controls, and do not apply extra effort to the task. The catecholaminergic system has been proposed to mediate the exertion of mental effort by reducing the noise of neuronal communication (Bouret & Richmond, 2015, Aston-Jones & Cohen, 2005; Servan-Schreiber et al., 1990; Walton & Bouret 2019). Either a reduced ability to keep information in working memory within prefrontal circuits, or to reduce the noise of cortical circuits by neuromodulators, would be related to a reduced wish to engage in difficult tasks (Westbrook & Braver, (2015); Manoach et al., 1999). This is also in agreement with the fact that negative symptoms often coexist with cognitive symptoms, and a lower threshold of cognitive demand would cause a feeling of helplessness. In sum, the complex interaction between cortical and neuromodulatory systems, rather than cortical circuits alone, may give rise to the observed WM impairments at high load.

Few fMRI studies have assessed activity in catecholaminergic areas during different behavioral conditions. Rausch et al. (2014) reported reduced activation in the VTA in medicated patients during a probabilistic decision-making task. Kohler et al. (2019) found reduced activation in the VTA in patients with schizophrenia during inhibitory cognitive control (Stroop task). Furthermore, activation in the VTA negatively correlated with negative symptoms in schizophrenia. LC activation in controls was correlated with RT, but not in patients. Cuervo-Lombard et al. (2012) reported decreased fMRI activation in the VTA in patients with schizophrenia during an autobiographical memory retrieval task. Together with these previous studies, our results of reduced activation in catecholaminergic nuclei associated to WM suggest reduced activation of the VTA as well as other catecholaminergic nuclei as a robust phenotype related to cognitive deficits in schizophrenia.

A few other studies using fMRI have assessed cortical activity during the N-back task. In general, reduced activity is reported for dorsal prefrontal areas with higher load (Carter et al. 1998; Glahn et al., 2005; Jiang et al., 2015; Gallucci et al., 2022), although this may be dependent on stimulus type (Brahmbhatt et al., 2006). A reduced activation of WM-related areas was suggested to reflect a disengagement of the task due to overload at high load levels (Thomas et al., 2021). Taken together, the reduced WM capacity observed in schizophrenia relays on general and specific effects across both cortical and subcortical systems, and the assessment or their joint function is necessary to understand the underlying mechanisms (Menon et al., 2023).

Several authors have proposed that cognitive symptoms in schizophrenia are partly due to a reduction of dopaminergic signaling on the prefrontal cortex. Supporting this view, recent multimodal studies found reduced release of DA on prefrontal areas (Fusar-Poli et al., 2011; Slifstein et al. 2015). An alteration of noradrenergic signaling on frontal targets has also been hypothesized (Arnsten, et al. 2012). Our results are consistent with a reduction of neuromodulatory action in cortical areas as a result of reduced activity in dopaminergic as well as noradrenergic structures. The dopaminergic and noradrenergic systems are tightly interconnected, and together facilitate higher order cognition (Chandler et al., 2014; Guiard et al., 2008; Harley, 2004; Smith & Greene, 2012; Vollbrecht, 2010; Xing et al., 2016). Importantly, these systems are highly flexible. Future work aiming to further clarify the joint function of catecholaminergic areas during cognition is necessary to clarify the mechanisms underlying cognitive deficits and possible remediation in schizophrenia.

Our study had several limitations. Contrary to cortical imaging, brainstem imaging is subjected to other sources of artifacts, and usually suffers from lower signal-to-noise ratio as copared to the cortex. Here we included an AROMA-based artifact detection method, which robustly deals with spurious effects (Pruim et al., 2015). In addition, we used a preprocessing pipeline that employed the ANTS tool for the coregistration step, thus ensuring proper coregistration of brainstem across subjects (Ewerts et al. 2019). Despite the relatively small sample size (N = 22 in the patient group), we detected significant activation in brainstem nuclei and were able to find a load-dependent difference between patients and controls. Another limitation was the fact that in the brainstem the structures are usually not so well defined as in the cortex. For example, the ventral tegmental area complex is composed of a heterogeneous population of dopaminergic and non-dopaminergic neurons. However, previous work using PET has shown that activity in the VTA correlates with dopamine release, and dopamine is involved in working memory processing, thus providing support to our findings of VTA involvement in the N-back task.

In conclusion, here we found that schizophrenia patients are comparable to controls when it comes to low working memory load, but at the highest load level, schizophrenia patients presented a reduced activation of catecholaminergic nuclei. This novel result highlights neuromodulatory activity as a possible signature of reduced cognitive function at higher load levels, and the importance of assessing activity in neuromodulatory regions together with the cortex. Importantly, neuromodulatory activity can be subjected to pharmacological interventions at different stages of the disease, offering possible avenues of remediation of cognitive symptoms.

## Notes

### Competing Interest Statement

The authors have declared no competing interest.

## References

1. Abi-Dargham, A., Rodenhiser, J., & Printz, D. (2000). Increased baseline occupancy of D2 receptors by dopamine in schizophrenia. PNAS, 97(14), 8104–8109.

2. Arnsten, A. F. T., Wang, M. J., & Paspalas, C. D. (2012). Neuromodulation of thought: Flexibilities and vulnerabilities in prefrontal cortical network synapses. Neuron, 76(1), 223–239.

3. Aston-Jones, G. & Cohen, J. D. (2005). An integrative theory of locus coeruleus-norepinephrine function: Adaptive gain and optimal performance. Annual Review of Neuroscience, 28, 403–450.

4. Barch, D. M. & Smith, E. (2008). The cognitive neuroscience of working memory: Relevance to CNTRICS and schizophrenia. Biological Psychiatry, 64(1), 11–17.

5. Beissner, F., Schumann, A., Brunn, F., Eisenträger, D., & Bär, K. (2014). Advances in functional magnetic resonance imaging of the human brainstem. Neuroimage, 86, 91–98.

6. Bianciardi, M., Toschi, N., Edlow, B. L., Eichner, C., Setsompop, K., Polimeni, J. R., Brown, E. N., Kinney, H. C., Rosen, B. R., & Wald, L. L. (2015). Toward an in vivo neuroimaging template of human brainstem nuclei of the ascending arousal, autonomic, and motor systems. Brain Connectivity, 5(10), 597–607.

7. Bouret, S. & Richmond, B. J. (2015). Sensitivity of locus ceruleus neurons to reward value for goal-directed actions. Journal of Neuroscience, 35(9), 4005–4014.

8. Brahmbhatt, S. B., Haut, K., Csernansky, J. G., & Barch, D. M. (2006). Neural correlates of verbal and nonverbal working memory deficits in individuals with schizophrenia and their high-risk siblings. Schizophrenia Research, 87(1-3), 191–204.

9. Brett, M., Anton, J. L., Valabregue, R., & Poline, J. B. (2002). Region of interest analysis using an SPM toolbox. Neuroimage, 13(2), 210–217.

10. Brooks, J. C. W., Faull, O. K., Pattinson, K. T. S., & Jenkinson, M. (2013). Psysiological noise in brainstem fMRI. Frontiers in Human Neuroscience, 7, Article 623.

11. Carter, C. S., Perlstein, W., Ganguli, R., Brar, J., Mintun, M., & Cohen, J. D. (1998). Functional hypofrontality and working memory dysfunction in schizophrenia. The American Journal of Psychiatry, 155(9), 1285–1287.

12. Chandler, D. J., Waterhouse, B. D., & Gao, W. (2014). New perspectives on catcholaminergic regulation of executive circuits: Evidence for independent modulation of prefrontal functions by midbrain dopaminergic and noradrenergic neurons. Frontiers in Neural Circuits, 8(53), Article 53.

13. Cardno, A. G. and Gottesman, I. I. (2000). Twin studies of schizophrenia: From bow-andarrow concordances to star wars Mx and functional genomics. American Journal of Medical Genetics, 97, (12-17).

14. Chen, A. P. F., Chen, L., Kim, T.A., & Xiong, Q. (2021). Integrating the roles of midbrain dopamine circuits in behavior and neuropsychiatric disease. Biomedicines, 9(6), 647.

15. Cuervo-Lombard, C., Lemogne, C., Gierski, F., Béra-Potelle, C., Tran, E., Portefaix, C., Kaladjian, A., Pierot, L., & Limosin, F. (2012). Neural basis of autobiographical memory retrieval in schizophrenia. British Journal of Psychiatry, 201(6), 473–480.

16. Deserno, L., Sterzer, P., Wüstenberg, T., Heinz, A., & Schlagenhauf, F. (2012). Reduced prefrontal-parietal effective connectivity and working memory deficits in schizophrenia. Journal of Neuroscience, 32(1), 12–20.

17. Esteban, O., Markiewicz, C.J., Blair, R.W. et al. (2019) fMRIPrep: a robust preprocessing pipeline for functional MRI. Nat Methods 16, 111–116.

18. Ewert, S., Horn, A., Finkel, F., Li, N., Kühn, A. A., & Herrington, T. M. (2019). Optimization and comparative evaluation of nonlinear deformation algorithms for atlas-based segmentation of DBS target nuclei. NeuroImage. 10.1016/j.neuroimage.2018.09.061

19. Fusar-Poli, F., Howes, O. D., Allen, P., Broome, M., Valli, I., Asselin, M., Montgomery, A. J., Grasby, P. M., & McGuire, P. (2011). Abnormal prefrontal activation directly related to pre-synaptic striatal dopamine dysfunction in people at clinical high risk for psychosis. Molecular Psychiatry, 16(1), 65–75.

20. Gallucci, J., Tan, T., Schifani, C., Dickie, E. W., Voineskos, A. N., & Hawco, C. (2022). Greater individual variability in functional brain activity during working memory performanc in Schizophrenia Spectrum Disorders (SDD). Schizophrenia Research, 248, 21–31.

21. Gjerde, P. F. (1983). Attentional capacity dysfunction and arousal in schizophrenia. Psychological Bulletin, 93(1), 57–72.

22. Glahn, D. C., Ragland, J. D., Abramoff, A., Barrett, J., Laird, A. R., Bearden, C. E., & Velligan, D. I. (2005). Beyond hypofrontality: A quantitative meta-analysis of functional neuroimaging studies of working memory in schizophrenia. Human Brain Mapping, 25(1), 60–69.

23. Granholm, E., Morris, S. K., Sarkin, A. J., Asarnow, R. F., & Jeste, D. V. (1997). Pupillary responses index overload of working memory resources in schizophrenia. Journal of Abnormal Psychology, 106(3), 458.

24. Guiard, B. P., El Mansari, M., Merali, Z., & Blier, P. (2008). Functional interactions between dopamine, serotonin and norepinephrine neurons: An in-vivo electrophysiological study in rats with monoaminergic lesions. International Journal of Neuropsychopharmacology, 11(5), 625–639.

25. Harley, C. W. (2004). Norepinephrine and dopamine as learning signals. Neural Plasticity, 11(3-4), 191–204.

26. Howes, O. D., Cummings, C., Chapman, G. E., & Shatalina, E. (2023). Neuroimaging in schizophrenia: An overview of findings and their implications for synaptic changes. Neuropsychopharmacology, 48, 151–167.

27. Jiang, S., Yan, H., Chen, Q., Tian, L., Lu, T., Tan, H., Yan, J., & Zhang, D. (2015). Cerebral inefficient activation in schizophrenia patients and their unaffected parents during the N-back working memory task: A family fMRI study. PloS One, 10(8), e0135468.

28. Kalkstein, S., Hurford, I., & Gur, R. C. (2010). Neurocognition in schizophrenia. Current Topic in Behavioral Neuroscience, 4, 373–390.

29. Köhler, S., Wagner, G., & Bär, K. (2019). Activation of brainstem and midbrain nuclei during cognitive control in medicated patients with schizophrenia. Human Brain Mapping, 40(1), 202–213.

30. Lee, J. & Park, S. (2005). Working memory impairments in schizophrenia: A meta-analysis. Journal of Abnormal Psychology, 114(4), 599–611.

31. Mäki-Marttunen, V., Andreassen, O. A., & Espeseth, T. (2020). The role of norepinephrine in the pathophysiology of schizophrenia. Neuroscience & Biobehavioral Reviews, 118, 298–314.

32. Mäki-Marttunen, V. & Espeseth, T. (2020). Uncovering the locus coeruleus: Comparison of localization methods for functional analysis. Neuroimage, 224, 117409.

33. Mäki-Marttunen, V., Hagen, T., & Espeseth, T. (2019). Task context load induces reactive cognitive control: An fMRI study on cortical and brain stem activity. *Cognitive, Affective*, & Behavioral Neuroscience, 19, 945–965.

34. Manoach, D. S., Press, D. Z., Thangaraj, V., Searl, M. M., Goff, D. C., Halpern, E., Saper, C. B., & Warach, S. (1999). Schizophrenic subjects activate dorsolateral prefrontal cortex during a working memory task, as measured by fMRI. Society of Biological Psychiatry, 45, 1128–1137.

35. Matt, E., Fischmeister, F. P. S., Amini, A., Robinson, S. D., Weber, A., Foki, T., Gizewski, E. R., & Beisteiner, R. (2019). Improving sensitivity, specificity, and reproducibility of individual brainstem activation. Brain Structure and Function, 224(8), 2823–2838.

36. McGrath, J., Saha, S., Chant, D., & Welham, J. (2008). Schizophrenia: A concise overview of incidence, prevalence, and mortality. Epidemiologic Reviews, 30, 67–76.

37. Menon, V., Palaniyappan, L., & Supekar, K. (2023). Integrative brain network and salience models of psychopathology and cognitive dysfunction in schizophrenia. Biological Psychiatry, 94(2), 108–120.

38. Motley, S. E. (2018). Relationship between neuromodulation and working memory in the prefrontal cortex: It’s complicated. Frontiers in neural circuits, 12, 31.

39. van Os, J. & Kapur, S. (2009). Schizophrenia. Lancet, 374(9690), 635–645.

40. Owen, M. J., Sawa, A., & Mortensen, P. B. (2016). Schizophrenia. Lancet, 388(10039), 86–97.

41. Perez-Costas, E., Melendez-Ferro, M., Robers, R. C. (2010). Basal ganglia pathology in schizophrenia: Dopamine connections and anomalies. Journal of Neurochemistry, 113, 287–302.

42. Pruim, R. H. R., Mennes, M., van Rooij, D., Llera, A., Buitelaar, J. K., & Beckmann, C. F. (2015). ICA-AROMA: A robust ICA-based strategy for removing motion artifacts from fMRI data. Neuroimage, 112, 267–277.

43. Rausch, V. H., Bauch, E. M., & Bunzeck, N. (2014). White noise improves learning by modulating activity in dopaminergic midbrain regions and right superior temporal sulcus. Journal of Cognitive Neuroscience, 26(7), 1469–1480.

44. Repovš, G., & Barch, D. (2012). Working memory related brain network connectivity in individuals with schizophrenia and their siblings. Frontiers in Human Neuroscience, 6, 1–15.

45. Rice, M. W., Roberts, R. C., Melendez-Ferro, M., & Perez-Costas, E. (2016). Mapping dopaminergic deficiencies in the substantia nigra/ventral tegmental area in schizophrenia. Brain Structure and Function, 221(1), 185–201.

46. Robbins, T. W. & Arnsten, A. F. T. (2009). The neuropsychopharmacology of fronto-executive function: Monoaminergic modulatio*n*. Annual Review of Neuroscience, 32, 267–287.

47. Schulz, J., Zimmermann, J., Sorg, C., Menegaux, A., & Brandl, F. (2022). Magnetic resonance imaging of the dopamine system in schizophrenia – A scoping review. Frontiers in Psychiatry, 13, 925476.

48. Sclocco, R., Beissner, F., Bianciardi, M., Polimeni, J. R., & Napadow, V. (2018). Challenges and opportunities for brainstem neuroimaging with ultrahigh field MRI. Neuroimage, 168, 412–426.

49. Servan-Schreiber, D., Printz, H., & Cohen, J. D. (1990). A network model of catecholamine effects: Gain, signal-to-noise ratio, and behavior. Science, 249(4971), 892–895.

50. Singh, K., Cauzzo, S., García-Gomar, M. G., Stauder, M., Vanello, N., Passino, C., & Bianciardi, M. (2022). Functional connectome of arousal and motor brainstem nuclei in living humans by 7 Tesla resting-state fMRI. NeuroImage, 249, 118865.

51. Slifstein, M., van de Giessen, E., Van Snellenberg, J., Thompson, J. L., Narendran, R., Gil, R., Hackett, E., Girgis, R., Ojeil, N., Moore, H., D’Souza, D., Malison, R. T., Huang, Y., Lim, K., Nabulsi, N., Carson, R. E., Lieberman, J. A., Abi-Dargham, A. (2015). Deficits in prefrontal cortical and extrastriatal dopamine release in schizophrenia: A positron emission tomographic functional magnetic resonance imaging study. JAMA Psychiatry, 72(4), 316–24.

52. Smith, C. C. & Greene, R. W. (2012). CNS dopamine transmission mediated by noradrenergic innervation. Journal of Neuroscience, 32(18), 6072–6080.

53. Stahl, S. M. (2018). Beyond the dopamine hypothesis of schizophrenia to three neural networks of psychosis: Dopamine, serotonin, and glutamate. CNS Spectrums, 23(3), 187–191.

54. Starc, M., Murray, J. D., Santamauro, N., Savic, A., Diehl, C., Cho, Y. T., Srihari, V., Morgan, P. T., Krystal, J. H., Wang, X., Repovs, G., & Anticevic, A. (2017). Schizophrenia is associatd with a pattern of spatial working memory deficits consistent with cortical disinhibition. Schizophrenia Research, 181, 107–116.

55. Sullivan, P. F., Kendler, K. S., Neale, M. C. (2003). Schizophrenia as a complex trait evidence from a meta-analysis of twin studies. Archives of General Psychiatry, 60(12), 1187–1192.

56. Thomas, M. L., Duffy, J. R., Swerdlow, N., Light, G. A., & Brown, G. G. (2022). Detecting the inverted-U in fMRI studies of schizophrenia: A comparison of three analysis methods. Journal of International Neuropsychological Society, 28(3), 258–269.

57. Tost, H., Alam, T., & Meyer-Lindenberg, A. (2010). Dopamine and psychosis: Theory, pathomechanisms and intermediate phenotypes. Neuroscience & Biobehavioral Reviews, 34(5), 689–700.

58. Turker, H. B., Riley, E., Luh, W., Colcombe, S. J., & Swallow, K. M. (2021). Estimates of locus coeruleus function with functional magnetic resonance imaging are influenced by localization approaches and the use of multi-echo data. Neuroimage, 236, 118047.

59. Van Kammen, D. P., & Kelley, M. (1991). Dopamine and norepinephrine activity in schizophrenia: An integrative perspective. Schizophrenia Research, 4(2), 173–191.

60. Varazzani, C., San-Galli, A., Gilardeau, S., & Bouret, S. (2015). Noradrenaline and dopamine neurons in the reward/efford trade-off: A direct electrophysiological comparison in behaving monkeys. Journal of Neuroscience, 35(20), 7866–7877

61. Volbrecht, P. (2010). Mechanisms for the interaction of dopamine and norepinephrine in the prefrontal cortex: Implications for the treatment of cognitive symptoms of schizophrenia. Vanderbilt Reviews Neuroscience, 2.

62. Walton, M. E. & Bouret, S. (2019). What is the relationship between dopamine and effort? Trends of Neuroscience, 42(2), 79–91.

63. Wang, M., Ramos, B. P., Paspalas, C. D., Shu, Y., Simen, A., Duque, A., Vijayraghavan, S., Brennan, A., Dudley, A., Nou, E., Mazer, J. A., McCormick, D. A., Arnsten, A. F. T. (2007). Alpha2A-adrenoceptors strengthen working memory networks by inhibiting cAMP-HCN channel signaling in prefrontal cortex. Cell, 129(2), 397–410.

64. Weinstein, J. J., Chohan, M. O., Slifstein, M., Kegeles, L. S., Moore, H., & Abi-Dargham, A. (2017). Biological Psychiatry, 81(1), 31–42.

65. Westbrook, A. & Braver, T.S. (2015). Cognitive effort: A neuroeconomic approach. Cognitive, Affective & Behavioral Neuroscience, 15(2), 395–415.

